# L-Lysine production from glucose and chitin monomers using engineered *Vibrio natriegens*

**DOI:** 10.64898/2026.04.10.717670

**Authors:** Elly Straube, Thi Van Anh Tran, Anna Faber, Nadine Ihle, Rubén Crespo Blanco, Ha Thanh Le, Georg Fritz, Cláudio J. R. Frazão, Thomas Walther

## Abstract

Despite its industrial importance, microbial L-lysine production has largely been confined to classical producer strains, leaving the fast-growing, non-pathogenic marine microorganism *V. natriegens* largely untapped as an unconventional biosynthetic platform. In this work, we established an L-lysine-overproducing *V. natriegens* DSM759 strain through a step-wise, systematic rational engineering strategy targeting the native biosynthetic pathway. Guided by our prior systems-level analysis of the strain’s genetic and regulatory architecture, we identified key metabolic bottlenecks and implemented knowledge-driven interventions to relieve pathway constraints. Central to production was alleviation of feedback inhibition in the native key regulatory enzymes, aspartate kinase (AK, *lysC*) and dihydrodipicolinate synthase (DHDPS, *dapA*). Site-directed amino-acid substitutions, replicating established *E. coli* feedback-resistance mechanisms, were introduced into conserved regions of the *V. natriegens* DSM759 enzymes, producing L-lysine-insensitive variants with kinetic parameters comparable to that of corresponding wild type enzymes. Among the tested configurations, the strain co-expressing *Vn.lysC2* and *Vn.dapA1:E84T* reached the highest L-lysine titer (9.0±0.6 mM) and yield (0.11±0.01 mol_Lys_ mol_Glc_^-1^), whereas overexpression of additional L-lysine pathway genes provided no further benefit. Leveraging the host’s metabolic versatility, L-lysine synthesis was also demonstrated from the chitin-derived amino-sugar N-acetylglucosamine (0.09±0.00 mol_Lys_ mol_GlcNAc_^-1^), highlighting the potential to valorize chitin-rich waste streams from the seafood industry. This work establishes a minimal, rational strategy for L-lysine biosynthesis in *V. natriegens* DSM759 and positions it as a promising platform for sustainable amino acid production.

## 1. Introduction

L-lysine is a basic, essential amino acid that plays a pivotal role in promoting growth and development, particularly in protein synthesis, hormone production, and immune function. Because animals cannot synthesize L-lysine *de novo* [1], this amino acid is a critical supplement in human nutrition and livestock feed [2]. Beyond its nutritional role, L-lysine is used in pharmaceutical and cosmetic formulations and serves as a precursor for bio-based chemicals such as putrescine, ε-poly-L-lysine, and nylon monomers [3–7]. To meet the increasing global demand, industrial L-lysine production relies predominantly on microbial fermentation using extensively engineered *Escherichia coli* and *Corynebacterium glutamicum* strains [3, 8]. Key strain engineering strategies include eliminating feedback inhibition of the rate-limiting enzymes and overexpressing biosynthetic genes in the L-lysine pathway [9, 10], optimising precursor and cofactor supply, and suppressing competing pathways. Systems-biology approaches, including genome-scale and enzyme-constrained modelling, have further guided the identification of pathway bottlenecks [11]. Industrial processes typically employ fed-batch fermentation to balance biomass formation with amino acid synthesis [12, 13], resulting in peak titers of 120-190 g L^-1^. Despite these advances, challenges remain, which include minimizing by-products, simplifying downstream purification, reducing substrate costs, and improving robustness in non-sterile conditions [3, 8, 14].

*Vibrio natriegens* has attracted increasing attention as a microbial production chassis due to its exceptionally high growth and substrate uptake rates, characteristics that offer the prospect of markedly shorter fermentation processes and improved space-time yields [15–17]. Over the past decade, its potential has been explored for the biosynthesis of diverse compounds, including organic and amino acids [18–20], biopolymers [21], pigments [22, 23], and short-chain alcohols [24, 25]. In parallel, a steadily expanding genetic toolbox – comprising standardized expression parts [26, 27] and multiple genome-editing methods [21, 28] – has significantly enhanced its tractability for metabolic engineering, positioning *V. natriegens* as a compelling platform for future biotechnological applications. Building on this foundation, we recently characterized the genomic organization and regulation of the L-lysine pathway in *V. natriegens*: As part of the aspartate-family amino acids, L-lysine is synthesized starting from L-aspartate through nine consecutive enzymatic steps *via* the diaminopimelate (DAP) pathway (Figure 1). L-lysine exerts regulatory control over its own biosynthesis at both the enzymatic and transcriptional levels, strongly inhibiting the activity of two key enzymes – mono-functional aspartate kinase (AK, Vn.LysC1) and dihydrodipicolinate synthase (DHDPS, Vn.DapA1) – while also repressing transcription of *Vn.lysC1* and *Vn.dapD*. The genetic organization and regulatory architecture of this pathway closely mirror those of *E. coli*, highlighting conserved mechanisms of feedback control. However, deviating from the pathway design found in *E. coli*, *V. natriegens* also encodes a second, intrinsically feedback-resistant mono-functional AK isozyme (Vn.LysC2), potentially providing a native bypass to one of the primary metabolic bottlenecks [29]. Nevertheless, the presence of dual-level regulation indicates that multiple control points must be addressed to achieve efficient L-lysine overproduction. Proof-of-concept studies using transcription factor-based biosensors have isolated L-lysine-secreting mutants producing up to 1 mM extracellularly, confirming pathway accessibility despite low initial titers [30]. Building on these findings, increased L-lysine biosynthesis in *V. natriegens* DSM759 was demonstrated by genomic integration of heterologous genes from *C. glutamicum*, confirming that production can be enhanced in this host through strain engineering [31]. However, the simultaneous introduction of multiple pathway components in the latter study precluded quantitative assessment of their individual impact on pathway flux and thus provided limited guidance for targeted flux optimization.

**Figure 1:**
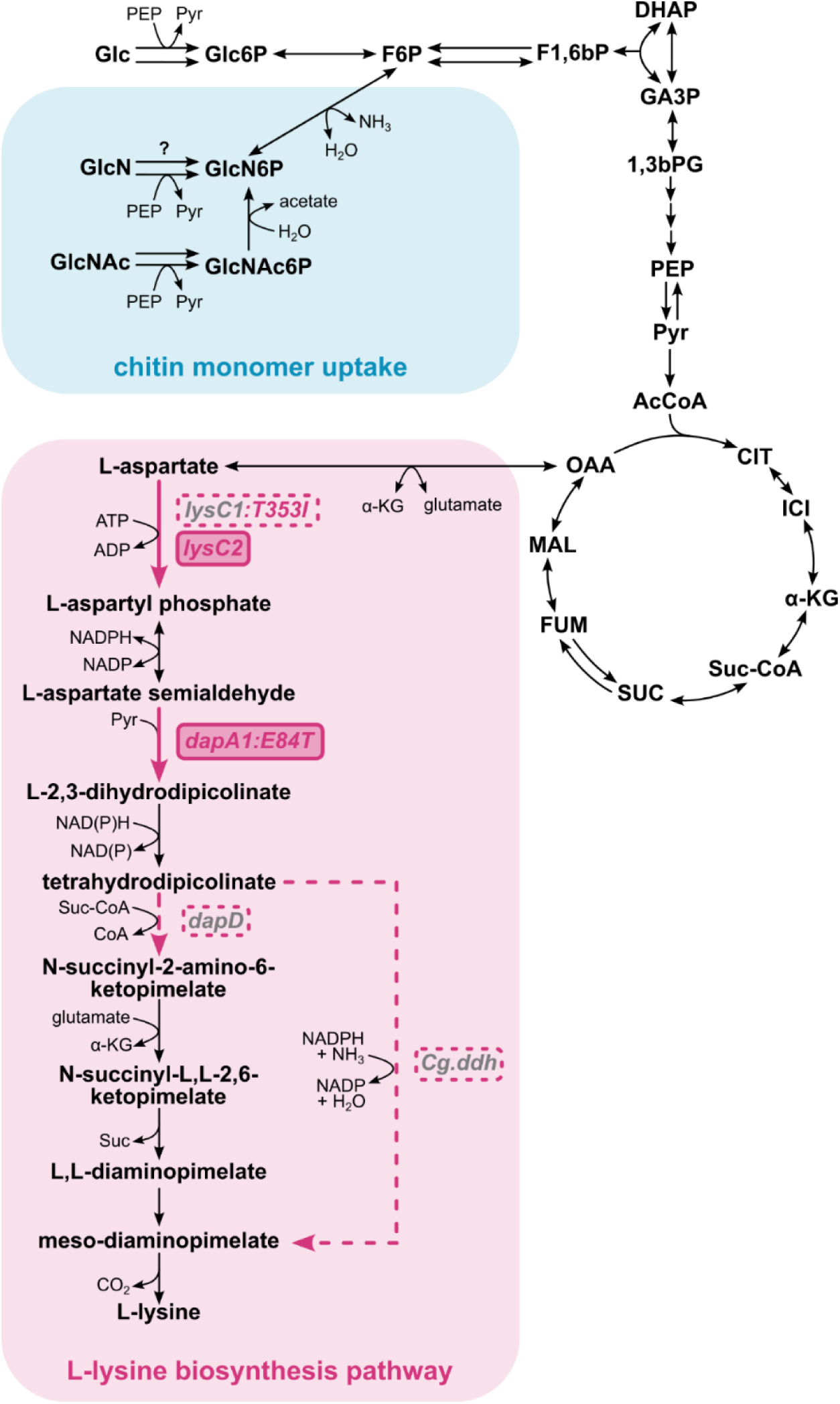
Biosynthesis of L-lysine from glucose and chitin monomers and applied engineering strategies for creation of a *V. natriegens* L-lysine overproducer strain. Glucose uptake, glycolysis and TCA cycle are shown in simplified form, while chitin monomer uptake and L-lysine biosynthesis pathways are depicted in detail, including side-products. Glucosamine (GlcN) import remains uncharacterized and is therefore indicated with a question mark. Tested plasmid-based (over-)expression strategies are indicated in pink dashed lines, whereas solid pink marks the overexpressed genes and reactions in the best-performing producer. Abbreviations: Glc – glucose; Glc6P – glucose-6-phosphate; F6P – fructose-6-phosphate; F1,6bP – fructose-1,6-bisphosphate; DHAP – dihydroxyacetone phosphate; GA3P – glyceraldehyde-3-phosphate; 1,3bPG – 1,3-bisphosphoglycerate; PEP – phosphoenolpyruvate; Pyr – pyruvate; AcCoA – acetyl-coenzyme A; CIT – citrate; ICI – isocitrate; α-KG – α-ketoglutarate; Suc-CoA – succinyl-coenzyme A; SUC – succinate; FUM – fumarate; MAL – malate; OAA – oxaloacetate. Gene names are written in grey, bold italic font. *lysC1(:T353I)* – gene encoding native L-lysine-sensitive, mono-functional aspartate kinase and engineered insensitive mutant; *lysC2* – gene encoding native L-lysine-insensitive, mono-functional aspartate kinase; *dapA1(:E84T)* – gene encoding native L-lysine-sensitive dihydrodipicolinate synthase and engineered insensitive mutant; *dapD –* gene encoding native diaminopimelate decarboxylase; *Cg.ddh* – gene encoding diaminopimelate dehydrogenase from *C. glutamicum*.

Here, we present a rational approach to engineer L-lysine production in *V. natriegens* DSM759 by optimizing the native biosynthetic pathway based on our prior pathway analysis. As a first step, feedback inhibition of key enzymes was alleviated by creating L-lysine-insensitive mutants of the native AK and DHDPS isozymes. The effects of these deregulated variants on L-lysine production were then systematically analysed through plasmid-based overexpression of the corresponding genes. To complement the alleviation of enzymatic regulation and further address potential rate-limiting steps, transcriptional repression of *Vn.dapD,* which encodes tetrahydrodipicolinate succinylase (DapD), was targeted. This was achieved by either plasmid-based overexpression of the native gene or by introduction of the heterologous diaminopimelate dehydrogenase (DapDH), which provides a bypass to the native reaction sequence. Finally, leveraging the metabolic versatility of *V. natriegens* DSM759, we examined L-lysine production from chitin-derived sugars as alternative feedstocks, illustrating the potential of this host to valorize chitin-rich crustacean shell waste. Together, these interventions establish a stepwise, systematic, rational framework for enabling L-lysine biosynthesis in *V. natriegens* DSM759 and highlight its promise as a fast-growing platform for sustainable amino-acid production.

## 2. Results and Discussion

### 2.1. Construction of L-lysine-resistant AK and DHDPS enzymes

The first step in the rational design of an L-lysine-producing *Vibrio natriegens* strain was to relieve feedback inhibition of the pathway’s key regulatory enzymes, AK and DHDPS. This strategy is well established for other microbial hosts, where native, L-lysine-sensitive enzymes support only minimal accumulation of this amino acid and significant L-lysine production typically only occurs after introducing feedback-resistant variants [9, 32–34]. Guided by these principles, we engineered the native *V. natriegens* enzymes through site-directed amino-acid substitutions whose positions were inferred from the known feedback-resistance mechanisms of the corresponding *E. coli* enzymes.

As the first target, we focused on AK, which catalyses the initial step of the L-lysine biosynthetic pathway, the phosphorylation of L-aspartate to L-aspartyl phosphate (Figure 1). In *V. natriegens*, two mono-functional AK isozymes have been characterized: Vn.LysC1, which is strongly inhibited by L-lysine, and Vn.LysC2, which is intrinsically L-lysine-insensitive [29]. Based on its regulatory profile and high sequence similarity to *E. coli* LysC, Vn.LysC1 was selected for mutagenesis. Allosteric inhibition of Ec.LysC by L-lysine occurs *via* binding at the ACT1 domain (residues 308-384). This triggers conformational changes that release the β15-αK loop, allowing rotation of the catalytic domain. These rearrangements block the ATP-binding site and shift the enzyme from the active R-state to the inactive T-state, providing the structural basis of inhibition [35]. Four amino-acid positions were previously reported to be crucial for L-lysine sensitivity in Ec.LysC (E250K [36], V339A [37], D340P [38], and T352I [36, 39]). They are fully conserved in Vn.LysC1 (E251K, V340A, D341P, and T353I) and were targeted for site-directed amino acid substitutions *via* inverse PCR. The L-lysine sensitivity of the resulting mutants was then evaluated *in vitro via* enzymatic assays, and compared to wild-type Vn.LysC1 and Vn.LysC2 activities (Figure 2): Baseline activities of the Vn.LysC1 variants, measured in the absence of L-lysine, showed that Vn.LysC1:V340A and Vn.LysC1:T353I exhibited elevated activities (115% and 152% compared to Vn.LysC1 wild-type), whereas Vn.LysC1:E251K and Vn.LysC1:D341P retained only 33-41% of the Vn.LysC1 wild-type activity. These observation are consistent with previous reports which showed that specific amino-acid substitutions at the regulatory site can affect basal enzyme efficiency even in the absence of L-lysine [10, 32, 37]. Under the same conditions, Vn.LysC2 reached 51% of Vn.LysC1 activity. Upon exposure to increasing L-lysine concentrations (2-100 mM), wild-type Vn.LysC1 displayed strong, concentration-dependent inhibition, retaining only 2-7% residual activity, while Vn.LysC2 remained largely insensitive, maintaining 44-56% residual activity across all tested concentrations. All four engineered Vn.LysC1 variants showed reduced sensitivity to L-lysine relative to the Vn.LysC1 wild-type enzyme, with the T353I variant exhibiting the most pronounced improvement, retaining 38% residual activity at 100 mM L-lysine. Kinetic characterization revealed that the T353I variant maintained a K_M_ value for L-aspartate comparable to wild-type Vn.LysC1 (4.3-5.4 mM) while achieving the highest v_max_ (183%), whereas Vn.LysC2 displayed the lowest v_max_ (53%) under the same conditions (Table 1). Based on this analysis, T353I was identified as the most promising engineered variant, while Vn.LysC2 remained of interest due to its intrinsic L-lysine resistance. Both enzymes were selected for further investigation.

**Figure 2:**
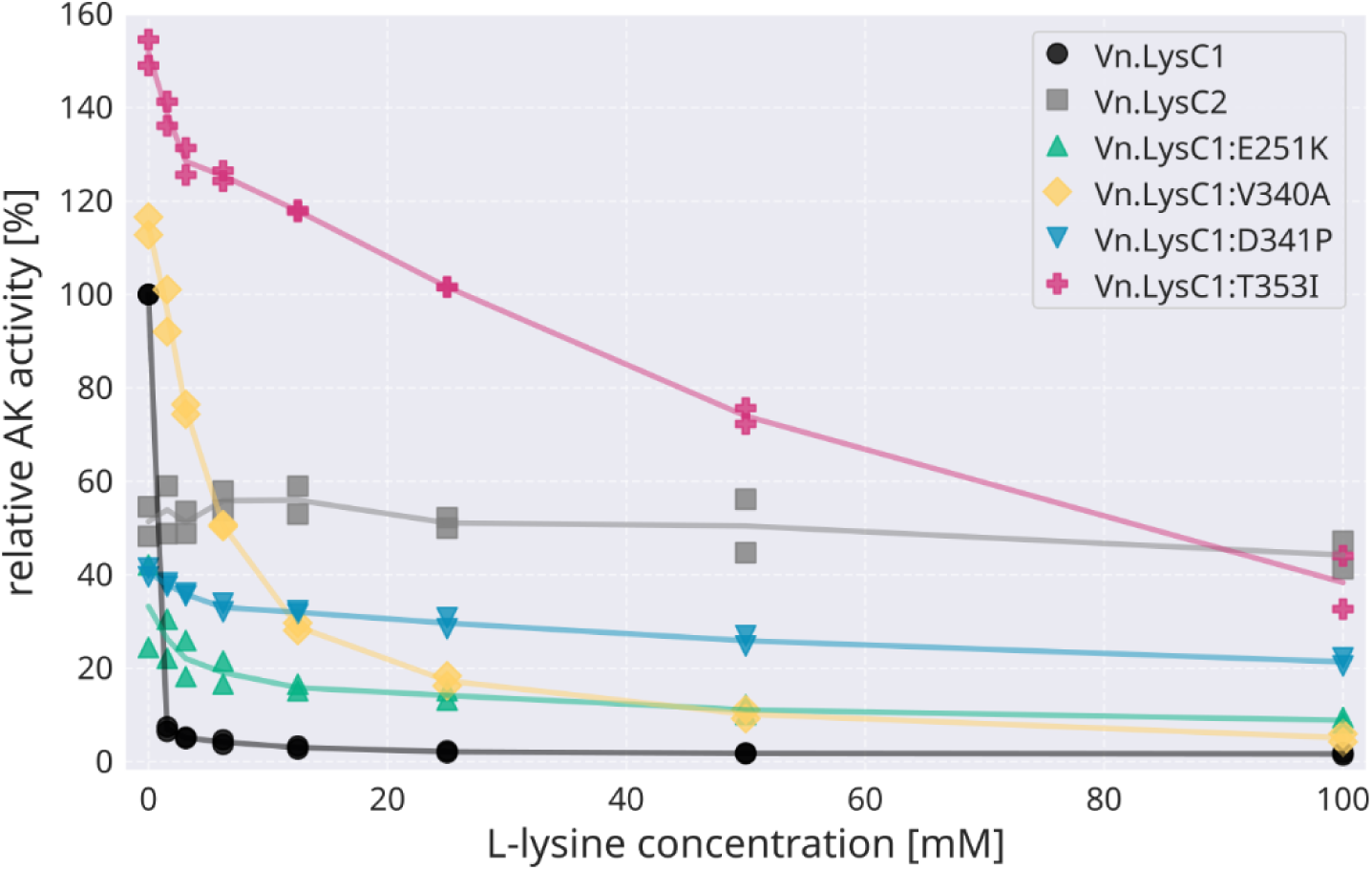
Construction of a L-lysine insensitive Vn.LysC1 mutant. The relative specific activities of native AK isozymes Vn.LysC1 and Vn.LysC2, and single amino acids mutants of Vn.LysC1, were determined with 50 mM L-aspartate as substrate in the presence of 0-100 mM L-lysine. Activities were normalized to the activity of Vn.LysC1 measured in the absence of L-lysine (v_max_ = 14.43 ± 2.81 U mg^-1^; pH 7.5; 37 °C; n = 2).

**Table 1.** Kinetic parameters of AK and DHDPS enzymes. ^a^.

| enzymatic activity | enzyme name | $v_{\max}$<br>[U mg<sup>-1</sup>] | $K_M$<br>[mM] | Reference |
| --- | --- | --- | --- | --- |
| AK<sup>b</sup> | Vn.LysC1 | 14.43±2.81 | 5.35±0.40 | [29] |
|  | Vn.LysC2 | 7.64±1.67 | 5.52±0.61 | [29] |
|  | Vn.LysC1:T353I | 26.46±1.80 | 4.29±0.87 | This study |
| DHDPS<sup>c</sup> | Vn.DapA1 | 19.18±4.20 | 0.20±0.01 | [29] |
|  | Vn.DapA1:E84T | 14.85±2.33 | 0.19±0.00 | This study |

<sup>a</sup> Experiments were carried out at pH 7.5 and 37 °C. Experimental data was fitted to the Michaelis-Menten model to determine kinetic parameters; n = 3.
<sup>b</sup> Kinetic parameters of AK activity were determined on 0.012-50 mM L-aspartate as substrate.
<sup>c</sup> Kinetic parameters of DHDPS activity were determined on 0.001-1 mM L-aspartate semialdehyde as substrate.

Subsequently, DHDPS was examined as the second regulatory enzyme of interest. DHDPS catalyses the condensation of L-aspartate semialdehyde (ASA) and pyruvate to form L-2,3-dihydrodipicolinate. The reaction is located at a key metabolic branch point since ASA also serves as a precursor for L-threonine, L-isoleucine, and L-methionine biosynthesis. In *V. natriegens*, two DHDPS isozymes have been identified, with Vn.DapA1 exhibiting approximately 270-fold higher specific activity than Vn.DapA2, indicating that Vn.DapA1 is the major functional isozyme [29]. Owing to its central metabolic role and strong inhibition by L-lysine, Vn.DapA1 was selected for engineering. Following the successful transfer of *E. coli* engineering strategies to *V. natriegens* in the case of AK, a single conserved position, E84, was chosen for mutagenesis. This residue corresponds to a well-characterized substitution in Ec.DapA that completely abolishes L-lysine inhibition while maintaining wild-type kinetic parameters [33]. Similarly, the resulting Vn.DapA1:E84T variant retained 86% residual activity in the presence of 5 mM L-lysine, compared to the strongly inhibited wild-type Vn.DapA1 (5% residual activity) (Figure 3). Thus, while feedback resistance was successfully conferred, the catalytic properties of the DHDPS enzyme were largely preserved, with kinetic parameters remaining in the same range (Table 1).

**Figure 3:**
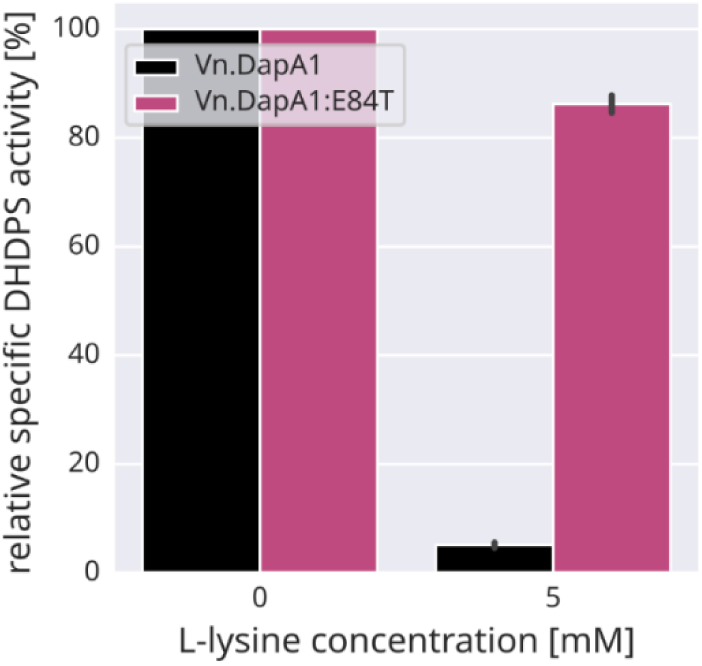
Construction of a L-lysine insensitive Vn.DapA1 mutant. The relative specific activities of native DHDPS Vn.DapA1 and single amino acids mutant Vn.DapA1:E84T were measured with 0.5 mM ASA as substrate in the presence of 0 and 5 mM L-lysine. Activities were normalized to the activity in the absence of L-lysine (pH 7.5; 37 °C; n = 3).

Overall, the engineering of L-lysine-resistant Vn.LysC1 and Vn.DapA1 variants was successful, demonstrating that strategies established in *E. coli* can be effectively transferred to *V. natriegens*. Their overexpression *in vivo* is expected to relieve key regulatory bottlenecks, providing a rational basis for the creation of L-lysine-overproducing *V. natriegens* strains, which is explored in the subsequent chapters.

### 2.2. Overexpression of L-lysine-resistant AK and DHDPS enzymes leads to L-lysine production

To assess the impact of L-lysine-resistant AK and DHDPS variants on *in vivo* L-lysine production, *V. natriegens* strains expressing the native or engineered enzymes were constructed. The genes were cloned into a medium-copy plasmid under the control of an inducible P_tac_ promoter, and expression was induced with IPTG (1 mM) in early exponential phase. For L-lysine production, cell growth is typically restricted to redirect carbon flux from biomass formation to product synthesis. In industrial settings, this is commonly achieved through fed-batch fermentation [2, 8], where carbon availability is carefully limited. Alternatively, auxotrophic growth factor limitation can be used, in which a specific amino acid – typically L-threonine – serves as the growth-limiting nutrient [12]. Here, for initial assessment of the enzyme variants’ impact on L-lysine production in shake flasks, growth was restricted using sulphur limitation. Restriction of sulphur availability induces stationary-phase conditions in the presence of excess carbon and nitrogen, which we considered favourable for L-lysine production. Producer strains were cultivated in 25 mL of sulphur-limited VN mineral medium containing 20 g L^-1^ glucose and 150 mM NH_4_Cl, buffered with 200 mM MOPS. L-lysine production was quantified after 24 h of cell cultivation (Figure 4).

**Figure 4:**
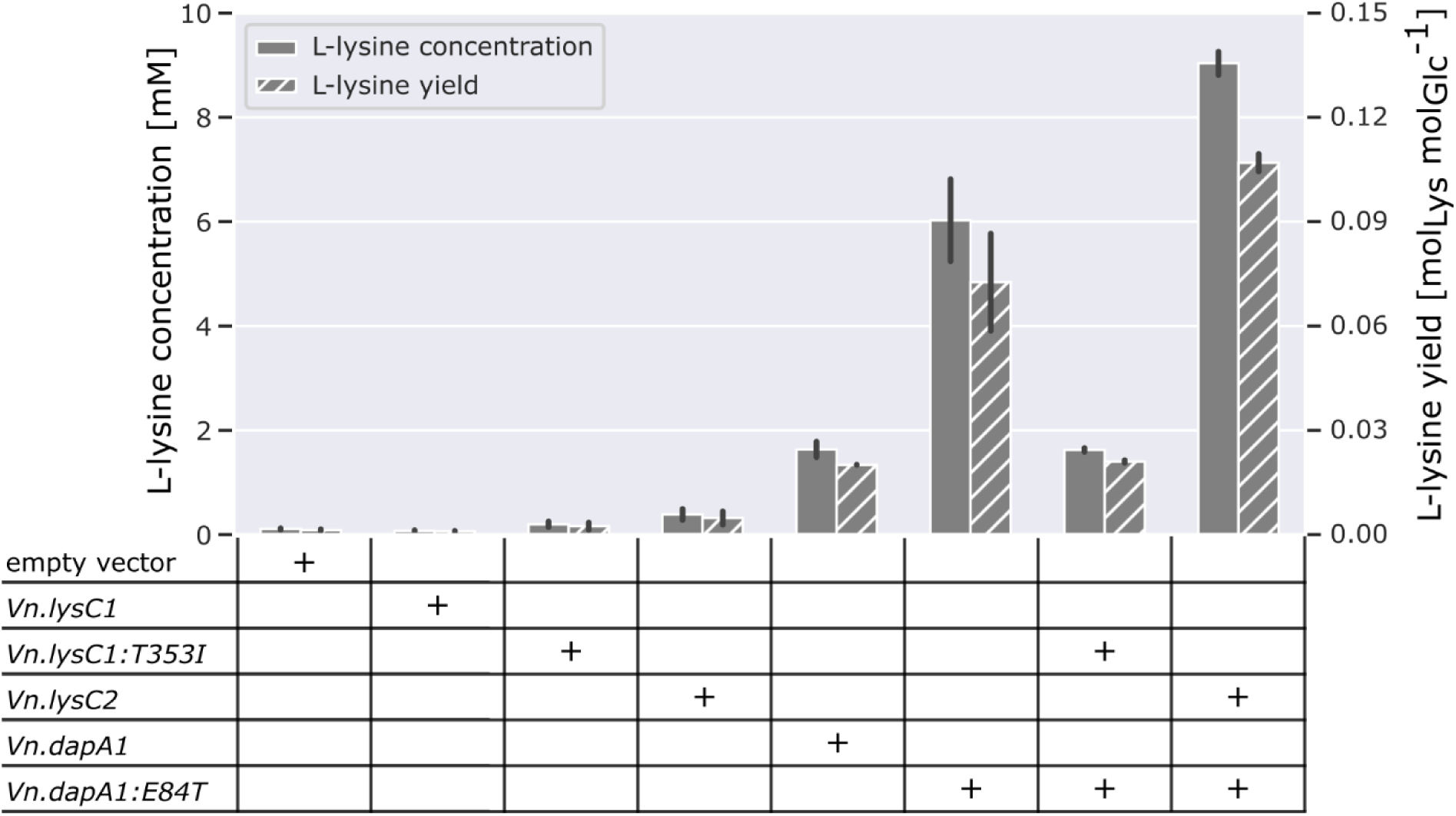
Plasmid-based overexpression of L-lysine-insensitive AK and DHDPS encoding genes increases L-lysine production in *V. natriegens*. L-lysine concentration after 24 h and L-lysine yield are shown for *V. natriegens* DSM759 *Δdns* strains harbouring different pMBI vectors during growth on glucose. The table beneath the plot indicates, with “+”, the gene variant expressed in each strain (S-limited VN mineral medium with 20 g L^-1^ glucose and 150 mM NH_4_Cl, 1 mM IPTG, 37 °C, 220 rpm, n≥2).

L-lysine production by the control strain harbouring an empty pMBI vector was negligible, reaching only 0.1±0.0 mM. Similarly, individual (over-)expression of AK variants resulted in none or only minor increases in L-lysine accumulation, with final concentrations of 0.1±0.0 mM (Vn.LysC1), 0.2±0.1 mM (Vn.LysC1:T353I), and 0.4±0.1 mM (Vn.LysC2), indicating that sole overexpression of AK variants was insufficient to substantially increase flux toward L-lysine. In contrast, individual overexpression of DHDPS variants led to markedly higher L-lysine production: overexpression of the wild-type *Vn.dapA1* gene resulted in formation of 1.6±0.1 mM L-lysine, while expression of the feedback-resistant *Vn.dapA1:E84T* mutant increased titers to 6.0±0.8 mM, consistent with DHDPS representing a major bottleneck in the L-lysine biosynthetic pathway. Co-expression of the L-lysine-insensitive AK and DHDPS variants revealed distinct effects depending on the AK enzyme used. Combination of the Vn.LysC1:T353I mutant with the Vn.DapA1:E84T mutant resulted in a pronounced reduction in L-lysine production (1.6±0.1 mM) compared to expression of the *Vn.dapA1:E84T* mutant alone. A possible explanation for this collapse is suboptimal expression of *Vn.lysC1:T353I*, potentially resulting from an unfavourable interaction between the upstream RBS and the *Vn.lysC1:T353I* coding sequence. In contrast, co-expression of the intrinsically L-lysine-resistant *Vn.lysC2* variant with the *Vn.dapA1:E84T* mutant yielded the highest L-lysine titers observed (9.0±0.6 mM), with a corresponding yield of 0.11±0.01 mol L-lysine per mol glucose. The strain predominantly produced L-lysine during the growth phase, prior to growth arrest induced by sulphur limitation (Supplementary Figure S1), indicating growth-associated production under the applied conditions. Combined with the observation that glucose was not fully consumed after 24 h, this highlights that medium composition and/or cultivation parameters could be further optimized to increase L-lysine production. Notably, this strain’s L-lysine levels exceeded the sum of the individual contributions of Vn.LysC2 and Vn.DapA1:E84T, suggesting a synergistic effect on pathway flux.

Overall, these results indicate that the DHDPS reaction represents a more significant control point in the L-lysine biosynthetic pathway than the AK reaction. A possible explanation was that native expression of *Vn.lysC2* is sufficient to sustain high pathway flux at the AK node, rendering DHDPS the primary limiting step. To test this hypothesis, L-lysine production was compared between the *Vn.lysC2* knockout and wild-type strains when only *Vn.dapA1:E84T* was overexpressed. Production remained comparable in both backgrounds (Supplementary Figure S3), suggesting that other native AK enzymes can compensate for *Vn.lysC2* deletion under these cultivation conditions. Vn.LysC1 is unlikely to assume this role due to its strong sensitivity to L-lysine inhibition. As *V. natriegens* encodes multiple functional AK enzymes, including bifunctional AK-HD variants, it remains unclear which enzyme(s) maintain flux at the AK node in the presence of L-lysine.

Taken together, these findings demonstrate that overexpression of feedback-resistant pathway enzymes is sufficient to trigger L-lysine accumulation without further strain modifications, consistent with rational design strategies reported for L-lysine-producing *E. coli* and *C. glutamicum* strains [9, 33]. Based on this foundation, subsequent analyses focused on the question whether expression of additional pathway genes further increases L-lysine production in *V. natriegens*.

### 2.3. Overexpression of additional L-lysine pathway enzymes does not further increase product yield

Following the alleviation of feedback inhibition of AK and DHDPS enzymes, we next aimed to fully relieve regulatory constraints along the L-lysine biosynthetic pathway. To this end, the *Vn.dapD* gene encoding tetrahydrodipicolinate succinylase (DapD), which is subject to transcriptional repression by L-lysine [29], was targeted for plasmid-based overexpression to circumvent this regulatory limitation. In parallel, expression of a heterologous diaminopimelate dehydrogenase (DapDH) from *C. glutamicum* (*Cg.ddh*) was evaluated as an alternative strategy to enhance flux by enabling direct conversion of tetrahydrodipicolinate to meso-diaminopimelate, thereby bypassing the native four-step succinylase route for L-lysine biosynthesis *via dapD* in *V. natriegens* (Figure 1).

To evaluate the impact of *Vn*.*dapD* or *Cg.ddh* (over-)expression, the respective genes were introduced separately into the best-performing producer strain (combined expression of genes encoding feedback-resistant Vn.LysC2 and Vn.DapA1:E84T enzymes). For heterologous *Cg.ddh*, both the wild-type gene and a manually codon-optimized variant (*Cg.ddh:CO*, Supplementary Text S1) were tested. Strains harbouring the respective plasmids were cultivated in sulphur-limited mineral medium with 20 g L^-1^ glucose as carbon source; gene expression was induced with 1 mM IPTG, and L-lysine accumulation was quantified after 24 h (Figure 5, Supplementary Figure S2).

**Figure 5:**
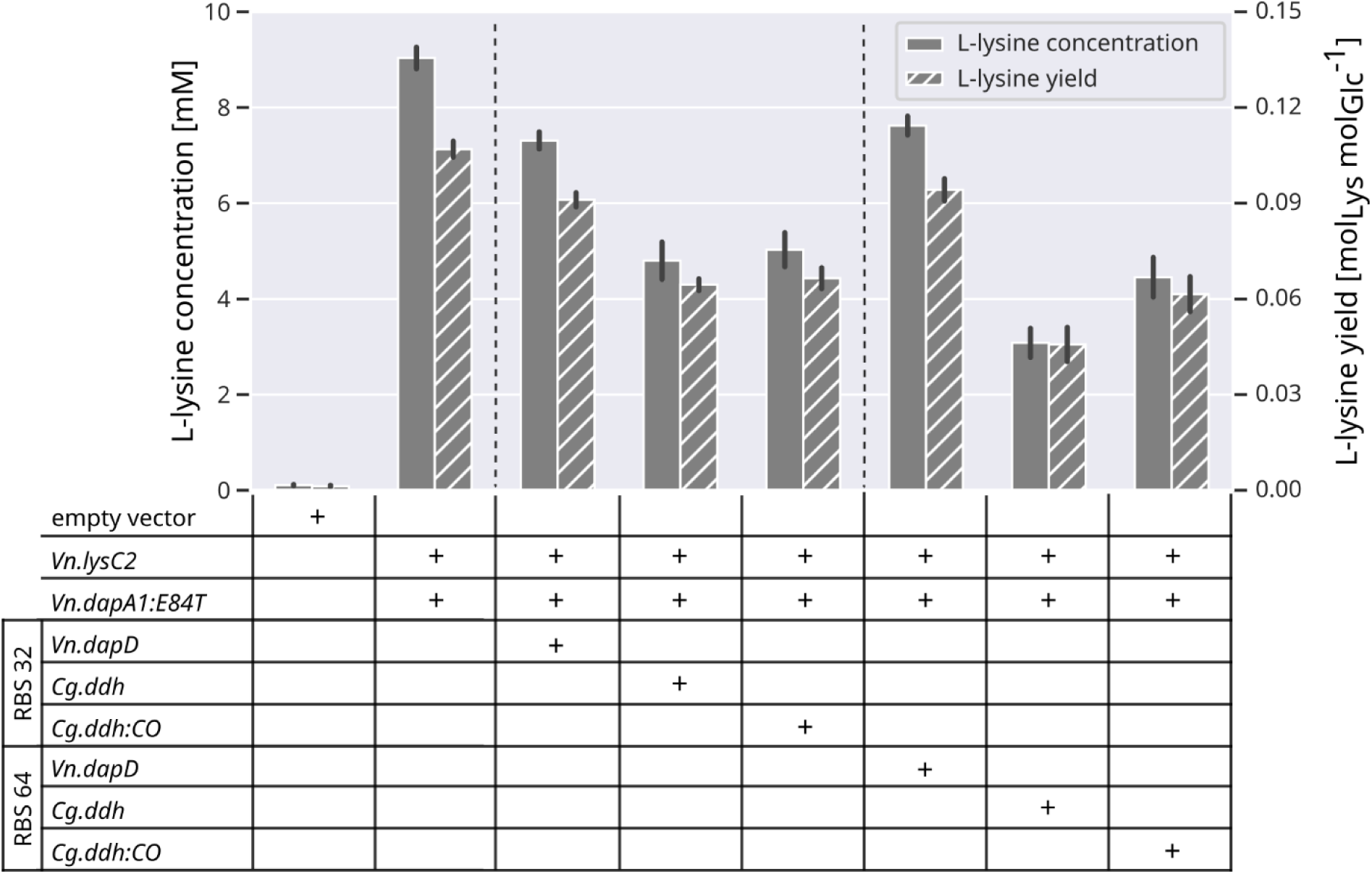
Effect of additional overexpression of native and heterologous pathway enzymes on L-lysine production. L-lysine concentration after 24 h and L-lysine yield are shown for *V. natriegens* DSM759 *Δdns* strains harbouring different pMBI vectors during growth on glucose. The table beneath the plot indicates, with “+”, the gene variant (and its RBS) expressed in each strain. (25 mL S-limited VN medium with 20 g L^-1^ glucose and 150 mM NH_4_Cl, 1 mM IPTG, 37°C, 220 rpm, n≥2).

We analysed the effect of expressing *Vn.dapD* and *Cg.ddh* using two RBS sequences of different strengths. Unexpectedly, additional plasmid-based overexpression of these genes did not enhance L-lysine production compared to the parent strain (expressing only *Vn.lysC2* and *Vn.dapA1:E84T*). When using a medium-strength RBS (RBS 32), overexpression of *Vn.dapD* caused a moderate reduction in L-lysine titers (to 81%), whereas heterologous expression of both wild-type and codon-optimized *Cg.ddh* resulted in a substantially stronger decrease (to 47% and 44%, respectively). Replacement of the RBS with a high-strength variant (RBS 64) did not restore L-lysine production: titers remained unchanged for *Vn.dapD* and codon-optimized *Cg.ddh:CO*, while expression of wild-type *Cg.ddh* further impaired L-lysine formation (reduction to 66%). While overexpression of *dapD* or *ddh* genes is an established strategy to improve L-lysine formation in *E. coli* and *C. glutamicum* [9, 10], our results indicate that simple amplification of this pathway segment is insufficient to increase L-lysine production in *V. natriegens* under the tested conditions. The observed production decline may result from suboptimal expression balance and the metabolic burden associated with plasmid-based overexpression. Additionally, the effectiveness of the heterologous dehydrogenase pathway may have been limited by physiological factors. Although the dehydrogenase route is energetically favourable, DapDH enzymes typically exhibit low affinity for ammonium, requiring elevated NH_4_^+^ concentrations to enhance L-lysine formation [40, 41]. While overexpression of *Cg.ddh* has shown mixed effects on L-lysine formation in *E. coli*, use of alternative DapDH from *Symbiobacterium thermophilum* was reported to be more compatible and beneficial [10, 40]. Together, these observations suggest that optimizing expression balance and ensuring enzyme-host compatibility may be required to further improve L-lysine production in *V. natriegens*.

### 2.4. L-lysine production using chitin monomers as carbon source

Chitin is the most abundant natural nitrogen-containing biopolymer and is widely found in crustacean shells, insect exoskeletons, and fungal cell walls. Large quantities of chitin-rich materials accumulate as low-value waste, particularly from seafood processing industries, making them an attractive and readily available feedstock for microbial production processes, especially for nitrogen-containing compounds such as amino acids, including L-lysine. Due to its insolubility and structural complexity, microbial utilization of chitin typically relies on prior hydrolysis to the soluble monomers N-acetyl-D-glucosamine (GlcNAc) and D-glucosamine (GlcN) [42, 43]. *V. natriegens* is particularly well suited for their utilization, as it exhibits exceptionally high uptake rates for both GlcNAc and GlcN (data not shown). To evaluate L-lysine production from these chitin-derived substrates, the best-performing producer strain expressing genes encoding feedback-resistant Vn.LysC2 and Vn.dapA1:E84T enzymes was cultivated in sulfur-limited mineral medium containing 100 mM GlcN or GlcNAc as the sole carbon source (Figure 6, Supplementary Figure S4).

**Figure 6:**
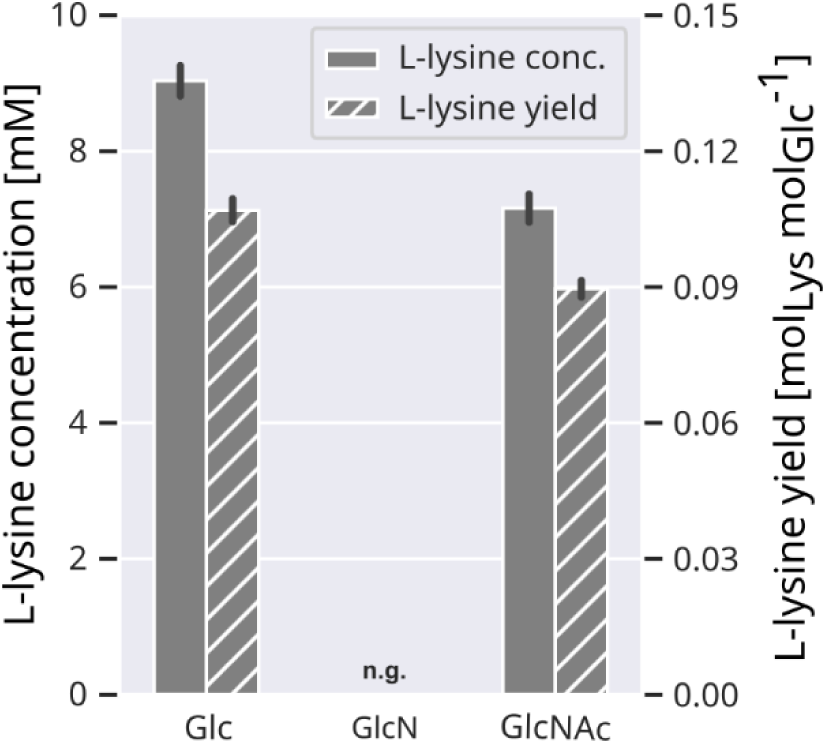
L-lysine production on chitin monomers as carbon source. L-lysine concentration after 24 h and L-lysine yield on respective carbon source are shown for *V. natriegens* DSM759 *Δdns* harboring the pMBI_Vn.lysC2_Vn.dapA1:E84T vector (25 mL S-limited VN medium with 100 mM C-source and 150 mM NH_4_Cl, 1 mM IPTG, 37°C, 220 rpm, n.g. – no growth, n≥3).

Cultivation on GlcNAc yielded L-lysine titers of 7.2±0.29 mM and a yield of 0.09±0.00 mol_Lys_ mol_GlcNAc_^-1^ after 24 h, comparable to those obtained on glucose, demonstrating efficient utilization of this chitin monomer for amino acid production. In contrast, no L-lysine formation was observed on GlcN, as the producer strain failed to grow under the applied cultivation conditions. This lack of growth likely reflects the combined effects of intrinsic limitations in GlcN metabolism and the additional metabolic burden imposed by plasmid maintenance. *E. coli* grows slower on GlcN than on GlcNAc, which has been attributed to differences in transport and regulatory induction of the shared GlcN/GlcNAc assimilation pathway (Figure 1) [44–46]. Wild-type *V. natriegens* DSM759 exhibits a similar pattern [20], suggesting that comparable regulatory constraints may restrict GlcN uptake and catabolism in the engineered host. The additional cellular demands associated with plasmid maintenance in the producer strain likely exacerbate these pre-existing limitations, ultimately preventing both growth and L-lysine formation on GlcN. Supplementation of the medium with yeast extract (0.5-2 g L^-1^) did not restore growth (data not shown), indicating that merely adding a richer nutrient source is insufficient to overcome these constraints. These findings highlight that enabling L-lysine production from GlcN will require targeted identification and engineering of the host’s chitin monomer uptake and assimilation pathways. This is particularly relevant because chemical hydrolysis of chitin frequently produces mixtures of GlcNAc and GlcN due to deacetylation [47]. In addition to strain engineering, co-feeding strategies – such as adding a small amount of GlcNAc to GlcN – may help induce the pathway and improve GlcN utilization [48].

In summary, L-lysine production from the chitin monomer GlcNAc was successfully demonstrated at levels comparable to glucose, highlighting the potential of the engineered strain for valorization of chitin-derived feedstocks. Achieving robust growth and L-lysine production on GlcN requires further metabolic and process optimization but represents an important step toward complete utilization of chitin hydrolysates in a sustainable biorefinery framework.

## 3. Conclusion

In this work, we constructed a L-lysine-overproducing *V. natriegens* DSM759 strain through a step-wise, rational engineering strategy targeting the native biosynthetic pathway. Guided by our prior systems-level analysis of the strain’s genetic and regulatory architecture, we identified key metabolic bottlenecks and implemented knowledge-driven interventions to relieve pathway constraints.

The first engineering step focused on enzymatic-level regulatory limitations, alleviating feedback inhibition of the key enzymes AK and DHDPS. Mutations known to confer L-lysine resistance in homologous *E. coli* enzymes were introduced into conserved regions of the native *V. natriegens* DSM759 enzymes, yielding L-lysine-insensitive variants with kinetic properties comparable to the wild-type enzymes. Systematic plasmid-based overexpression of these native and engineered variants revealed that the combination of Vn.LysC2 and Vn.DapA1:E84T resulted in the best-performing strain, achieving a titer of 9.0±0.6 mM L-lysine at a yield of 0.11±0.01 mol_Lys_ mol_Glc_^-1^ and enabling initial L-lysine accumulation without further strain modifications (Figure 1). To further increase pathway flux, regulatory limitations at the transcriptional level were addressed through plasmid-based overexpression of native *Vn.dapD* and, alternatively, heterologous *Cg.ddh*. These interventions did not further enhance L-lysine production; instead, titers and yields declined, underscoring the importance of balanced pathway expression and the need for additional strain optimization. Exploiting the host’s feedstock flexibility, L-lysine production from GlcNAc was achieved at levels comparable to glucose utilization. In contrast, the engineered strain was unable to grow on GlcN, despite robust growth of the wild type, revealing limitations in substrate assimilation under conditions of increased metabolic demand.

Overall, the best strain, which overexpressed only *Vn.lysC2* and *Vn.dapA1:E84T*, required minimal engineering yet achieved yields comparable to the most advanced reported *V. natriegens* DSM759 L-lysine producer strain that relies on extensive heterologous pathway expression and gene deletions (0.15 mol_Lys_ mol_Glc_^-1^) [31]. These findings demonstrate that a knowledge-driven optimization of the native pathway is effective to unlock substantial L-lysine production capacity. Further improvements will require both additional strain engineering and cultivation optimization. On the genetic level, fine-tuning expression of key genes – *via* RBS libraries or chromosomal integration of engineered variants – could enhance stability and industrial relevance, while engineering robust GlcN utilization may broaden substrate scope. In parallel, process optimization appears essential: L-lysine formation occurred predominantly during the growth phase prior to sulfur limitation, and incomplete glucose consumption suggested non-optimal conditions. Medium refinement, fed-batch strategies, and increased ammonium concentrations to favor flux through DapDH enzyme represent promising directions.

In summary, we constructed a minimally engineered yet efficient L-lysine-producing *V. natriegens* DSM759 strain through rational, step-wise optimization of its native pathway, providing a solid foundation for future metabolic and process engineering toward industrial application.

## 4. Materials and Methods

### 4.1. Reagents and chemicals

Chemicals, solvents and oligonucleotides were purchased from Sigma-Aldrich (Darmstadt, Germany), unless stated otherwise. Plasmid DNA purification and gel extraction kits as well as restriction enzymes were purchased from NEB (Frankfurt am Main, Germany). Sanger sequencing was carried out by Eurofins (Ebersberg, Germany).

### 4.2. Strains and plasmids

All strains and plasmids used in this study are listed in Tables 2 and 3.

**Table 2.** *Escherichia coli* and *Vibrio natriegens* strains used in this study.

| Strain | Genotype | Reference/Origin |
| --- | --- | --- |
| NEB® 5-alpha | *E. coli fhuA2::IS2 Δ(mmuP-mhpD)169 ΔphoA8 glnX44 φ80d[ΔlacZ58(M15)] rfbD1 gyrA96 luxS11 recA1 endA1 rphWT thiE1 hsdR17* | NEB™ |
| NEB® stable | *E. coli F' proA<sup>+</sup>B<sup>+</sup> lacI<sup>q</sup> Δ(lacZ)M15 zzf::Tn10 (Tet<sup>R</sup>)/Δ(ara-leu) 7697 araD139 fhuA ΔlacX74 galK16 galE15 e14- Φ80dlacZΔM15 recA1 relA1 endA1 nupG rpsL (Str<sup>R</sup>) rph spoT1 Δ(mrr-hsdRMS-mcrBC)* | NEB™ |
| BL21 (DE3) | *E. coli fhuA2 [lon] ompT gal (λ DE3) [dcm] ΔhsdS* | NEB™ |
| *V. natriegens* | *V. natriegens* DSM759 (ATCC14048) Δ*dns* | [26] |

**Table 3.** Plasmids used in this study.

| Plasmid | Relevant characteristics | Reference /Origin |
| --- | --- | --- |
| *In vitro studies* |  |  |
| pET28a(+) | *f1* origin, Kan<sup>R</sup>, T7 promoter | Novagen™ |
| pET28-Vn.lysC1 | pET28a(+) derivative with N-terminal his-tagged *Vn.lysC1* | [29] |
| pET28-Vn.lysC2 | pET28a(+) derivative with N-terminal his-tagged *Vn.lysC2* | [29] |
| pET28-Vn.lysC1:E251K | pET28a(+) derivative with N-terminal his-tagged *Vn.lysC1:E351K* | This study |
| pET28-Vn.lysC1:V340A | pET28a(+) derivative with N-terminal his-tagged *Vn.lysC1:V340A* | This study |
| pET28-Vn.lysC1:D341P | pET28a(+) derivative with N-terminal his-tagged *Vn.lysC1:D341P* | This study |
| pET28-Vn.lysC1:T353I | pET28a(+) derivative with N-terminal his-tagged *Vn.lysC1:T353I* | This study |
| pET28-Vn.dapA1 | pET28a(+) derivative with N-terminal his-tagged *Vn.dapA1* | [29] |
| pET28-Vn.dapA1:E84T | pET28a(+) derivative with N-terminal his-tagged *Vn.dapA1:E84T* | This study |
| *In vivo L-lysine production* |  |  |
| pMBI-GFP dropout | pMBI origin, Kan<sup>R</sup>(Vn), *lacI* under $P_{LacI}$, transcriptional unit: $P_{tac}$ – RBS (B0032) – GFP dropout – terminator (B0015) | This study |
| pMBI | empty pMBI derivative (without GFP dropout) | This study |
| pMBI_Vn.lysC1 | pMBI derivative carrying *Vn.lysC1* | This study |
| pMBI_Vn.lysC1:T353I | pMBI derivative carrying *Vn.lysC1:T353I* | This study |
| pMBI_Vn.lysC2 | pMBI derivative carrying *Vn.lysC2* | This study |
| pMBI_Vn.dapA1 | pMBI derivative carrying *Vn.dapA1* | This study |
| pMBI_Vn.dapA1:E84T | pMBI derivative carrying *Vn.dapA1:E84T* | This study |
| pMBI_Vn.lysC1:T353I_Vn.dapA1:E84T | pMBI derivative carrying *Vn.lysC1:T353I* and *Vn.dapA1:E84T* | This study |
| pMBI_Vn.lysC2_Vn.dapA1:E84T | pMBI derivative carrying *Vn.lysC2* and *Vn.dapA1:E84T* | This study |
| pMBI_Vn.lysC2_Vn.dapA1:E84T_Vn.dapD(RBS32) | pMBI derivative carrying *Vn.lysC2*, *Vn.dapA1:E84T* and *Vn.dapD* (*Vn.dapD* preceded by RBS B0032) | This study |
| pMBI_Vn.lysC2_Vn.dapA1:E84T_Vn.dapD(RBS64) | pMBI derivative carrying *Vn.lysC2*, *Vn.dapA1:E84T* and *Vn.dapD* (*Vn.dapD* preceded by RBS B0064) | This study |
| pMBI_Vn.lysC2_Vn.dapA1:E84T_Cg.ddh(RBS32) | pMBI derivative carrying *Vn.lysC2*, *Vn.dapA1:E84T* and *Cg.ddh* (*Cg.ddh* preceded by RBS B0032) | This study |
| pMBI_Vn.lysC2_Vn.dapA1:E84T_Cg.ddh(RBS64) | pMBI derivative carrying *Vn.lysC2*, *Vn.dapA1:E84T* and *Cg.ddh* (*Cg.ddh* preceded by RBS B0064) | This study |
| pMBI_Vn.lysC2_Vn.dapA1:E84T_Cg.ddh:CO(RBS32) | pMBI derivative carrying *Vn.lysC2*, *Vn.dapA1:E84T* and codon-optimised variant of *Cg.ddh* (*Cg.ddh:CO* preceded by RBS B0032) | This study |
| pMBI_Vn.lysC2_Vn.dapA1:E84T_Cg.ddh:CO(RBS64) | pMBI derivative carrying *Vn.lysC2*, *Vn.dapA1:E84T* and codon-optimised variant of *Cg.ddh* (*Cg.ddh:CO* preceded by RBS B0064) | This study |

### 4.3. Media

LB medium (LB Broth (Luria/Miller), Carl Roth) and LB agar plates (LB agar (Luria/Miller), Carl Roth) were used for cloning procedures, protein production and recovery of cells from glycerol stocks (25% v/v, stored at -80 °C). For work with *V. natriegens*, LB medium was supplemented with v2 salts (204 mM NaCl, 4.2 mM KCl and 23.14 mM MgCl_2_; LBv2) [49]. Where required, kanamycin sulfate was added at 50 mg L^-1^ for *E. coli* and 200 mg L^-1^ for *V. natriegens*.

For *in vivo* L-lysine production in shake-flask experiments, an adapted sulfur-limited VN mineral medium was used [20], containing 100 mM of carbon source (glucose, glucosamine, or N-acetyl- glucosamine), 150 mM NH_4_Cl, 15 g L^-1^ NaCl, 1 g L^-1^ KH_2_PO_4_, 1 g L^-1^ K_2_HPO_4_, 2 mM MgCl_2_, 10 mg L^-1^ CaCl_2_, 16.4 mg L^-1^ FeSO_4_·7 H_2_O and trace elements (10 mg L^-1^ MnSO_4_·H_2_O, 0.3 mg L^-1^ CuSO_4_·5 H_2_O, 1 mg L^-1^ ZnSO_4_·7 H_2_O and 0.02 mg L^-1^ NiCl_2_·6 H_2_O). The media was buffered with 200 mM MOPS.

### 4.4. Side-directed mutagenesis, gene expression and protein purification for *in vitro* enzyme studies

Site-directed mutagenesis to generate L-lysine-resistant AK and DHDPS isozymes was performed by inverse PCR [50] using the primer pairs listed in Table S1. Each reaction contained 6 ng of the pET28 plasmid carrying the respective wild-type enzyme as template. PCR was carried out with Q5 polymerase (NEB), using a reduced cycle number of 18. Residual template DNA was removed by DpnI digestion (NEB), and the resulting plasmids were transformed into chemically competent NEB® 5-alpha (NEB) and verified by Sanger sequencing. All constructed plasmids are listed in Table 3.

Gene expression and protein purification were carried out as previously described by Straube *et al*., 2026 [29].

### 4.5. AK and DHDPS enzymatic assay

Enzymatic assays to determine kinetic parameters and L-lysine resistance of wild-type and mutant AK and DHDPS enzymes were performed as previously described by Straube *et al*., 2026 [29].

### 4.6. Plasmid construction for *in vivo* L-lysine production

All plasmids constructed and used for *in vivo* L-lysine biosynthesis are listed in Table 3. In this study, we designed and assembled the pMBI plasmid backbone, which contains a GFP dropout cassette serving as a placeholder for insertion of target genes within a single transcriptional unit (pMBI-GFP dropout). The backbone was assembled using Golden Gate cloning following the strategy of Stukenberg *et al.*, 2021 [26].

Expression constructs were generated by replacing the GFP dropout cassette with one, two, or three coding sequences along with their respective ribosome binding sites (RBSs). RBSs were assigned according to gene position within the operon – B0032 (first gene), B0033 (second), and B0032/B0064 (third) – and were sourced from the Vnat Collection [26, 27]. Constructs were assembled using isothermal assembly (NEBuilder® HiFi DNA Assembly Master Mix, NEB) following the manufacturer’s instructions. Target genes were amplified by PCR from the respective pET28 plasmids or from *V. natriegens* DSM759 chromosomal DNA using Q5 polymerase (NEB), with primers listed in Table S1. The pMBI-GFP dropout backbone was linearized by BbsI digestion (NEB), and fragments were purified by gel extraction (Monarch® Spin DNA Gel Extraction Kit, NEB) prior to assembly.

An empty pMBI vector was generated as a control by PCR amplification of the digested backbone to create overlapping regions, followed by self-ligation using NEBuilder® HiFi DNA Assembly Master Mix.

Plasmids carrying the *Cg.ddh* gene and/or RBS B0032 upstream of the third gene of the transcriptional unit were generated by restriction-ligation-based cloning. The backbone (pMBI_Vn.lysC2 _Vn.dapA1:E84T) and the *Vn.dapD*/*Cg.ddh* inserts were amplified by PCR using Q5 polymerase. Wild-type *Cg.ddh* was amplified from *Corynebacterium glutamicum* DSM20300 chromosomal DNA, and the codon-optimized variant was obtained from a synthetic gene (Azenta). PCR primers introduced NdeI and EcoRI (for insertion of *Vn.dapD*) or NdeI and SalI restriction sites (for insertion of *Cg.ddh*) at the 5′ and 3′ ends (Table S1). Both backbone and inserts were digested with NdeI and EcoRI/SalI (NEB), purified by gel extraction, and ligated using T4 DNA ligase (NEB) according to the manufacturer’s protocol.

All constructed pMBI vectors were transformed into chemically competent NEB® 5-alpha or NEB® stable (NEB) and verified by Sanger sequencing.

### 4.7. Shake-flask cultivation for L-lysine production

Shake-flask cultivations were performed at 30 or 37 °C, 220 rpm in an orbital shaker (Ecotron, Infors). For each experiment, *V. natriegens* cells were freshly transformed with the respective pMBI plasmids following the protocol of Stukenberg *et al.*, 2021 [26]. For precultures, single colonies were inoculated into 5 mL LBv2 medium in 100 mL shake flasks and incubated overnight at 30 °C. Cells were harvested by centrifugation (4500 g, 10 min, room temperature), washed with 9 g L^-1^ NaCl, and resuspended to inoculate the main cultures at a starting OD_600 nm_ of 0.2. Main cultures were grown in 25 mL sulfur-limited VN mineral medium in 250 mL baffled shake flasks at 37 °C. When cultures reached an OD_600 nm_ of 0.6–0.8, gene expression was induced by addition of 1 mM IPTG. Samples were collected at regular intervals, centrifuged (13000 g, 5 min, room temperature), and the supernatant stored at -20 °C until further analysis.

### 4.8. Analytical methods

Substrate (glucose, N-acetyl glucosamine) and metabolite concentrations (acetate, pyruvate, and L-lysine) were quantified by HPLC. Supernatants (1 mL) were filtered through 0.22 µm regenerated cellulose filters (VWR) and transferred into 1.5 mL HPLC vials (VWR). Samples were analysed on an AZURA® 862 bar HPLC system (Knauer), equipped with a P 6.1 L pump (LPG), AS 2.1 L autosampler (set to 6 °C), CT 2.1 column oven, RI detector (RID 2.1 L; set to 45 °C), and UV/Vis detector (MWD 2.1 L). The injection volume was 20 µL.

Sugars and organic acids were separated on a Rezex^TM^ ROA-Organic Acid H^+^ (8%) column (300 × 7.8 mm; Phenomenex) with a SecurityGuard^TM^ Carbo-H^+^ precolumn (4 × 3 mm; Phenomenex). Elution was performed with 0.5 mM H_2_SO_4_ at 0.5 mL min^-1^, and the column temperature was maintained at 80 °C.

L-lysine was quantified using a Chirex 3126 (D)-penicillamine column (150 × 4.6 mm; Phenomenex) protected by a SecurityGuard^TM^ AQ C18 cartridge (4 × 3 mm; Phenomenex). Elution was performed with 2 mM CuSO_4_ as mobile phase at 0.5 mL min^-1^, and the column temperature was maintained at 25 °C. To ensure the removal of late-eluting or otherwise retained compounds and to maintain long-term column performance, each chromatographic run included a methanol ramp in the mobile phase up to 5 % (v/v), followed by a return to 0 %.

### 4.9. Computational methods

Pairwise local sequence alignments were performed to determine whether amino acid residues in *V. natriegens* AK and DHDPS corresponding to L-lysine-resistant positions in *E. coli* were conserved. Amino acid sequences of the *E. coli* wild-type enzymes and the respective *V. natriegens* homologs were obtained from the KEGG database [51] and aligned using the EMBOSS Water algorithm (Smith-Waterman) *via* the EMBOSS web server [52]. Default parameters were used.

The sequence of *Corynebacterium glutamicum* diaminopimelate dehydrogenase *(Cg.ddh)* was manually codon-optimized for expression in *V. natriegens*. Rare codons throughout the gene were identified and optimized using the Graphical Codon Usage Analyser (GCUA; gcua.schoedl.de) based on the *V. natriegens* codon-usage table from Lee *et al.,* 2019 [53]. Additionally, the first 27 bases of the sequence were analysed using RNAfold (ViennaRNA package) [54] to predict potential secondary structures, and adjusted as needed to minimize stable base-pairing. The final codon-optimized gene was synthesized by Azenta, Inc., and the nucleotide sequence is provided in the Supplementary Material (Supplementary Text S1).

## Acknowledgements

This study was supported by a grant of the Federal Ministry of Research and Education (grant number 031B1076) awarded to TW.

## References

1. Lopez MJ, and Mohiuddin SS (2025). Biochemistry, Essential Amino Acids. StatPearls Publishing, Treasure Island (FL).

2. Cheng J, Chen P, Song A, Wang D, and Wang Q (2018). Expanding lysine industry: industrial biomanufacturing of lysine and its derivatives. J Ind Microbiol Biotechnol. 45(8): 719–734. doi: 10.1007/s10295-018-2030-8.

3. Wu Z, Chen T, Sun W, Chen Y, and Ying H (2024). Optimizing Escherichia coli strains and fermentation processes for enhanced L-lysine production: a review. Front Microbiol. 15. doi: 10.3389/fmicb.2024.1485624.

4. Li S, Mao Y, Zhang L, Wang M, Meng J, Liu X, Bai Y, and Guo Y (2022). Recent advances in microbial ε-poly-L-lysine fermentation and its diverse applications. Biotechnol Biofuels Bioprod. 15(1): 65. doi: 10.1186/s13068-022-02166-2.

5. Xu Z, Xu Z, Feng X, Xu D, Liang J, and Xu H (2016). Recent advances in the biotechnological production of microbial poly(ɛ-l-lysine) and understanding of its biosynthetic mechanism. Appl Microbiol Biotechnol. 100(15): 6619–6630. doi: 10.1007/s00253-016-7677-3.

6. Jorge JMP, Pérez-García F, and Wendisch VF (2017). A new metabolic route for the fermentative production of 5-aminovalerate from glucose and alternative carbon sources. Bioresour Technol. 245: 1701–1709. doi: 10.1016/j.biortech.2017.04.108.

7. Kumar R, Shah S, Paramita Das P, Bhagavanbhai GGK, Al Fatesh A, and Chowdhury B (2019). An overview of caprolactam synthesis. Catal Rev. 61(4): 516–594. doi: 10.1080/01614940.2019.1650876.

8. Liu J, Xu J-Z, Rao Z-M, and Zhang W-G (2022). Industrial production of L-lysine in *Corynebacterium glutamicum*: Progress and prospects. Microbiol Res. 262: 127101. doi: 10.1016/j.micres.2022.127101.

9. Becker J, Zelder O, Häfner S, Schröder H, and Wittmann C (2011). From zero to hero—Design-based systems metabolic engineering of *Corynebacterium glutamicum* for l-lysine production. Metab Eng. 13(2): 159–168. doi: 10.1016/j.ymben.2011.01.003.

10. Xu J, Han M, Ren X, and Zhang W (2016). Modification of aspartokinase III and dihydrodipicolinate synthetase increases the production of l-lysine in *Escherichia coli*. Biochem Eng J. 114: 79–86. doi: 10.1016/j.bej.2016.06.025.

11. Ye C, Luo Q, Guo L, Gao C, Xu N, Zhang L, Liu L, and Chen X (2020). Improving lysine production through construction of an Escherichia coli enzyme-constrained model. Biotechnol Bioeng. 117(11): 3533–3544. doi: 10.1002/bit.27485.

12. Kiss RD, and Stephanopoulos G (1991). Metabolic activity control of the L-lysine fermentation by restrained growth fed-batch strategies. Biotechnol Prog. 7(6): 501–509. doi: 10.1021/bp00012a004.

13. Ying H, He X, Li Y, Chen K, and Ouyang P (2014). Optimization of Culture Conditions for Enhanced Lysine Production Using Engineered Escherichia coli. Appl Biochem Biotechnol. 172(8): 3835–3843. doi: 10.1007/s12010-014-0820-7.

14. Yin L, Zhou Y, Ding N, and Fang Y (2024). Recent Advances in Metabolic Engineering for the Biosynthesis of Phosphoenol Pyruvate–Oxaloacetate–Pyruvate-Derived Amino Acids. Molecules. 29(12): 2893. doi: 10.3390/molecules29122893.

15. Hoff J, Daniel B, Stukenberg D, Thuronyi BW, Waldminghaus T, and Fritz G (2020). Vibrio natriegens: an ultrafast-growing marine bacterium as emerging synthetic biology chassis. Environ Microbiol. 22(10): 4394–4408. doi: 10.1111/1462-2920.15128.

16. Thoma F, and Blombach B (2021). Metabolic engineering of Vibrio natriegens. Essays Biochem. 65(2): 381–392. doi: 10.1042/EBC20200135.

17. Hädrich M, Schulze C, Hoff J, and Blombach B Vibrio natriegens: Application of a Fast-Growing Halophilic Bacterium. Springer, Berlin, Heidelberg; pp 1–32.

18. Wu F, Wang S, Peng Y, Guo Y, and Wang Q (2023). Metabolic engineering of fast-growing Vibrio natriegens for efficient pyruvate production. Microb Cell Factories. 22(1): 172. doi: 10.1186/s12934-023-02185-0.

19. Thoma F, Schulze C, Gutierrez-Coto C, Hädrich M, Huber J, Gunkel C, Thoma R, and Blombach B (2022). Metabolic engineering of Vibrio natriegens for anaerobic succinate production. Microb Biotechnol. 15(6): 1671–1684. doi: 10.1111/1751-7915.13983.

20. Hoffart E, Grenz S, Lange J, Nitschel R, Müller F, Schwentner A, Feith A, Lenfers-Lücker M, Takors R, and Blombach B (2017). High Substrate Uptake Rates Empower Vibrio natriegens as Production Host for Industrial Biotechnology. Appl Environ Microbiol. 83(22): e01614–17. doi: 10.1128/AEM.01614-17.

21. Dalia TN, Hayes CA, Stolyar S, Marx CJ, McKinlay JB, and Dalia AB (2017). Multiplex Genome Editing by Natural Transformation (MuGENT) for Synthetic Biology in Vibrio natriegens. ACS Synth Biol. 6(9): 1650–1655. doi: 10.1021/acssynbio.7b00116.

22. Wang Z, Tschirhart T, Schultzhaus Z, Kelly EE, Chen A, Oh E, Nag O, Glaser ER, Kim E, Lloyd PF, Charles PT, Li W, Leary D, Compton J, Phillips DA, Dhinojwala A, Payne GF, and Vora GJ (2020). Melanin Produced by the Fast-Growing Marine Bacterium Vibrio natriegens through Heterologous Biosynthesis: Characterization and Application. Appl Environ Microbiol. 86(5): e02749–19. doi: 10.1128/AEM.02749-19.

23. Ellis GA, Tschirhart T, Spangler J, Walper SA, Medintz IL, and Vora GJ (2019). Exploiting the Feedstock Flexibility of the Emergent Synthetic Biology Chassis Vibrio natriegens for Engineered Natural Product Production. Mar Drugs. 17(12): 679. doi: 10.3390/md17120679.

24. Erian AM, Freitag P, Gibisch M, and Pflügl S (2020). High rate 2,3-butanediol production with *Vibrio natriegens*. Bioresour Technol Rep. 10: 100408. doi: 10.1016/j.biteb.2020.100408.

25. Zhang Y, Li Z, Liu Y, Cen X, Liu D, and Chen Z (2021). Systems metabolic engineering of *Vibrio natriegens* for the production of 1,3-propanediol. Metab Eng. 65: 52–65. doi: 10.1016/j.ymben.2021.03.008.

26. Stukenberg D, Hensel T, Hoff J, Daniel B, Inckemann R, Tedeschi JN, Nousch F, and Fritz G (2021). The Marburg Collection: A Golden Gate DNA Assembly Framework for Synthetic Biology Applications in Vibrio natriegens. ACS Synth Biol. 10(8): 1904–1919. doi: 10.1021/acssynbio.1c00126.

27. Faber A, Politan RJ, Stukenberg D, Morris KM, Kim R, Jeon E, Inckemann R, Becker A, Thuronyi B, and Fritz G (2025). Expanding genetic engineering capabilities in Vibrio natriegens with the Vnat Collection. Nucleic Acids Res. 53(13): gkaf580. doi: 10.1093/nar/gkaf580.

28. Stukenberg D, Hoff J, Faber A, and Becker A (2022). NT-CRISPR, combining natural transformation and CRISPR-Cas9 counterselection for markerless and scarless genome editing in Vibrio natriegens. Commun Biol. 5(1): 1–13. doi: 10.1038/s42003-022-03150-0.

29. Straube E, Radde J, Tran TVA, Yazdi NK, Blanco RC, Le HT, Frazão CJR, and Walther T (2026). Genetic make-up and regulation of the L-lysine biosynthesis pathway in Vibrio natriegens. Microb Cell. 13: 44–62. doi: 10.15698/mic2026.02.867.

30. Stella RG, Baumann P, Lorke S, Münstermann F, Wirtz A, Wiechert J, Marienhagen J, and Frunzke J (2021). Biosensor-based isolation of amino acid-producing *Vibrio natriegens* strains. Metab Eng Commun. 13: e00187. doi: 10.1016/j.mec.2021.e00187.

31. Nghiem TT, Tran HL, Nguyen TT, Le TH, and Le QH (2025). Unlocking the potential of Vibrio natriegens for L-lysine production. Res J Biotechnol. 11(20): 1. doi: 10.25303/2011rjbt0109.

32. Chen Z, Meyer W, Rappert S, Sun J, and Zeng A-P (2011). Coevolutionary Analysis Enabled Rational Deregulation of Allosteric Enzyme Inhibition in Corynebacterium glutamicum for Lysine Production ▿. Appl Environ Microbiol. 77(13): 4352–4360. doi: 10.1128/AEM.02912-10.

33. Geng F, Chen Z, Zheng P, Sun J, and Zeng A-P (2013). Exploring the allosteric mechanism of dihydrodipicolinate synthase by reverse engineering of the allosteric inhibitor binding sites and its application for lysine production. Appl Microbiol Biotechnol. 97(5): 1963–1971. doi: 10.1007/s00253-012-4062-8.

34. Ohnishi J, Mitsuhashi S, Hayashi M, Ando S, Yokoi H, Ochiai K, and Ikeda M (2002). A novel methodology employing Corynebacterium glutamicum genome information to generate a new L-lysine-producing mutant. Appl Microbiol Biotechnol. 58(2): 217–223. doi: 10.1007/s00253-001-0883-6.

35. Kotaka M, Ren J, Lockyer M, Hawkins AR, and Stammers DK (2006). Structures of R- and T-state *Escherichia coli* Aspartokinase III. J Biol Chem. 281(42): 31544–31552. doi: 10.1016/S0021-9258(19)84068-1.

36. Kikuchi Y, Kojima H, and Tanaka T (1999). Mutational analysis of the feedback sites of lysine-sensitive aspartokinase of Escherichia coli. FEMS Microbiol Lett. 173(1): 211–215. doi: 10.1111/j.1574-6968.1999.tb13504.x.

37. Chen Z, Rappert S, Sun J, and Zeng A-P (2011). Integrating molecular dynamics and co-evolutionary analysis for reliable target prediction and deregulation of the allosteric inhibition of aspartokinase for amino acid production. J Biotechnol. 154(4): 248–254. doi: 10.1016/j.jbiotec.2011.05.005.

38. Wang Y, Li Q, Zheng P, Guo Y, Wang L, Zhang T, Sun J, and Ma Y (2016). Evolving the l-lysine high-producing strain of Escherichia coli using a newly developed high-throughput screening method. J Ind Microbiol Biotechnol. 43: 1227–1235. doi: 10.1007/s10295-016-1803-1.

39. Ogawa-Miyata Y, Kojima H, and Sano K (2001). Mutation Analysis of the Feedback Inhibition Site of Aspartokinase III of Escherichia coli K-12 and its Use in L-Threonine Production. Biosci Biotechnol Biochem. 65(5): 1149–1154. doi: 10.1271/bbb.65.1149.

40. Xu J-Z, Ruan H-Z, Liu L-M, Wang L-P, and Zhang W-G (2019). Overexpression of thermostable meso-diaminopimelate dehydrogenase to redirect diaminopimelate pathway for increasing L-lysine production in Escherichia coli. Sci Rep. 9: 2423. doi: 10.1038/s41598-018-37974-w.

41. Liu N, Zhang T-T, Rao Z-M, Zhang W-G, and Xu J-Z (2021). Reconstruction of the Diaminopimelic Acid Pathway to Promote L-lysine Production in Corynebacterium glutamicum. Int J Mol Sci. 22(16): 9065. doi: 10.3390/ijms22169065.

42. Ma X, Gözaydın G, Yang H, Ning W, Han X, Poon NY, Liang H, Yan N, and Zhou K (2020). Upcycling chitin-containing waste into organonitrogen chemicals via an integrated process. Proc Natl Acad Sci. 117(14): 7719–7728. doi: 10.1073/pnas.1919862117.

43. Abidin MZ, Junqueira-Gonçalves MP, Khutoryanskiy VV, and Niranjan K (2017). Intensifying chitin hydrolysis by adjunct treatments – an overview. J Chem Technol Biotechnol. 92(11): 2787–2798. doi: 10.1002/jctb.5208.

44. Álvarez-Añorve LI, Calcagno ML, and Plumbridge J (2005). Why Does Escherichia coli Grow More Slowly on Glucosamine than on N-Acetylglucosamine? Effects of Enzyme Levels and Allosteric Activation of GlcN6P Deaminase (NagB) on Growth Rates. J Bacteriol. 187(9): 2974–2982. doi: 10.1128/jb.187.9.2974-2982.2005.

45. Álvarez-Añorve LI, Bustos-Jaimes I, Calcagno ML, and Plumbridge J (2009). Allosteric Regulation of Glucosamine-6-Phosphate Deaminase (NagB) and Growth of Escherichia coli on Glucosamine. J Bacteriol. 191(20): 6401–6407. doi: 10.1128/jb.00633-09.

46. Álvarez-Añorve LI, Gaugué I, Link H, Marcos-Viquez J, Díaz-Jiménez DM, Zonszein S, Bustos-Jaimes I, Schmitz-Afonso I, Calcagno ML, and Plumbridge J (2016). Allosteric Activation of Escherichia coli Glucosamine-6-Phosphate Deaminase (NagB) In Vivo Justified by Intracellular Amino Sugar Metabolite Concentrations. J Bacteriol. 198(11): 1610–1620. doi: 10.1128/jb.00870-15.

47. Einbu A, and Vårum KM (2008). Characterization of Chitin and Its Hydrolysis to GlcNAc and GlcN. Biomacromolecules. 9(7): 1870–1875. doi: 10.1021/bm8001123.

48. Le T, Nguyen T-H, Vu D-C, Cao T-H-T, Vu OT-K, Nguyen T-T, Pham T-A, and Le T-H (2026). High-cell-density cultivation of Vibrio natriegens N5.3 on chitin monomers: a step toward chitin valorization. Biotechnol Lett. 48(1): 24. doi: 10.1007/s10529-026-03696-7.

49. Weinstock MT, Hesek ED, Wilson CM, and Gibson DG (2016). Vibrio natriegens as a fast-growing host for molecular biology. Nat Methods. 13(10): 849–851. doi: 10.1038/nmeth.3970.

50. Zheng L, Baumann U, and Reymond J-L (2004). An efficient one-step site-directed and site-saturation mutagenesis protocol. Nucleic Acids Res. 32(14): e115. doi: 10.1093/nar/gnh110.

51. Kanehisa M, and Goto S (2000). KEGG: Kyoto Encyclopedia of Genes and Genomes. Nucleic Acids Res. 28(1): 27–30. doi: 10.1093/nar/28.1.27.

52. Madeira F, Madhusoodanan N, Lee J, Eusebi A, Niewielska A, Tivey ARN, Lopez R, and Butcher S (2024). The EMBL-EBI Job Dispatcher sequence analysis tools framework in 2024. Nucleic Acids Res. 52(W1): W521–W525. doi: 10.1093/nar/gkae241.

53. Lee HH, Ostrov N, Wong BG, Gold MA, Khalil AS, and Church GM (2019). Functional genomics of the rapidly replicating bacterium Vibrio natriegens by CRISPRi. Nat Microbiol. 4(7): 1105–1113. doi: 10.1038/s41564-019-0423-8.

54. Gruber AR, Lorenz R, Bernhart SH, Neuböck R, and Hofacker IL (2008). The Vienna RNA Websuite. Nucleic Acids Res. 36(suppl_2): W70–W74. doi: 10.1093/nar/gkn188.

