## Supplementary Data for "L-Lysine production from glucose and chitin monomers using engineered *Vibrio natriegens*"

**Text S1| Codon-optimised *Cg.ddh* derivative.** Manually optimised codons are highlighted in red.

atgaccaacatccgtgtagctatcgtgggctacggaaacctgggacgcagcgtcgaaaagcttattgccaagcagccagacatggaccttgtag
gaatcttctcgcgccgtgccaccctcgacacaaagacgccagtctttgatgtcgccgacgtggacaagcacgccgacgacgtggacgtgctgttc
ctgtgcatgggctccgccaccgacatccctgagcaggcaccaaagttcgcgcagttcgcctgcaccgtagacacctacgacaaccaccgcgacat
cccacgccaccgccaggtcatgaacgaagccgccaccgcagccggcaacgttgcactggtctctaccggctgggatccaggaatgttctccatca
accgcgtctacgcagcggcagtcttagccgagcaccagcagcacaccttctggggcccaggtttgtcacagggccactccgatgctttgcgacgc
atccctggcgttcaaaaggcagtccagtacaccctcccatccgaagacgccctggaaaaggcccgccgcggcgaagccggcgaccttaccgga
aagcaaacccacaagcgccaatgcttcgtggttgccgacgcggccgatcacgagcgcatcgaaaacgacatccgcaccatgcctgattacttcgt
tggctacgaagtcgaagtcaacttcatcgacgaagcaaccttcgactccgagcacaccggcatgccacacggtggccacgtgattaccaccggc
gacaccggtggcttcaaccacaccgtggaatacatcctcaagctggaccgaaacccagatttcaccgcttcctcacagatcgctttcggtcgcgca
gctcaccgcatgaagcagcagggccaaagcggagctttcaccgtcctcgaagttgctccatacctgctctccccagagaacttggacgatctgat
cgcacgcgacgtctaa

**Table S1| Primers used in this study.**

| Primer | Sequence (5'→3') |
|---|---|
| *Primers for site-directed mutagenesis (SDM)* | |
| SDM-Vn.lysC1:E251K-fw | gaagcgtcaAAAatggcaaacttcggtgc |
| SDM-Vn.lysC1:E251K-rv | gcaccgaagtttgccatTTTgacgcttc |
| SDM-Vn.lysC1:V340A-fw | GCAgacctgattaccacttcagaaat |
| SDM-Vn.lysC1:V340A-rv | TGCtgaaattttgtgtttagccagaatctcg |
| SDM-Vn.lysC1:D341P-fw | CCActgattaccacttcagaaatcag |
| SDM-Vn.lysC1:D341P-rv | TGGcactgaaattttgtgtttagcca |
| SDM-Vn.lysC1:T353I-fw | gtttcgttaATCctagaccaaacagac |
| SDM-Vn.lysC1:T353I-rv | tttggtctagGATtaacgaaacactgatttc |
| SDM-Vn.dapA1:E84T-fw | cgggtgcaaatgctacccacACCgccgtgacgttcag |
| SDM-Vn.dapA1:E84T-rv | agtttactgaacgtcacggcGGTgtgggtagcatttgcac |
| *Construction of pMBI vectors via isothermal assembly (IA) or restriction-ligation-based cloning (RL)* | |
| IA-bb-fw | agtcacacaggaaagtaatcgcttaaccaggcatcaaataaaac |
| IA-bb-rv | gattactttcctgtgtgactc |
| IA-Vn.lysC1(bb)-fw | agtcacacaggaaagtaatcaATGAGCGCATTTAACGTAG |
| IA-Vn.lysC1-rv | TTATTTTTCAAATAGCTCAGC |
| IA-Vn.lysC1(bb)-rv | gttttatttgatgcctggttATTTTTCAAATAGCTCAGCATG |
| IA-Vn.lysC2(bb)-fw | agtcacacaggaaagtaatcaATGAAAAAGCCCCTTATCG |
| IA-Vn.lysC2-rv | TCACCCTATTAGGGCATG |
| IA-Vn.lysC2(bb)-rv | gttttatttgatgcctggttACCCTATTAGGGCATGTTTTTG |
| IA-Vn.dapA1(bb)-fw | agtcacacaggaaagtaatcaATGTTTTCAGGAAGTATCG |
| IA-Vn.dapA1(Vn.lysC1)-fw | ctgagctatttgaaaaataatactagagtcacacaggactaatcaATGTTTTCAGGAAGTATCG |
| IA-Vn.dapA1(Vn.lysC2)-fw | aacatgccctaatagggtgatactagagtcacacaggactaatcaATGTTTTCAGGAAGTATCG |
| IA-Vn.dapA1-rv | TTAATCTTTATAAATACGGGCG |
| IA-Vn.dapA1(bb)-rv | gttttatttgatgcctggttAATCTTTATAAATACGGGCG |
| IA-Vn.dapD-fw | cccgtatttataaagattaatactagagaaagaggggaaataatcaATGGCTTTCTTTTCTCTAG |
| IA-Vn.dapD-rv | gttttatttgatgcctggttAGTTGTTTGAGTGTAGTTC |
| RL-bb-fw | attcgtgtcgaCCAGGCATCAAATAAAACG |
| RL-bb-rv | tagactcatatgttaatctttataaatacgggcgtc |
| RL-Cg.ddh-fw | attcgtcatatgTACTAGAGAAAGAGGGGAAATAATCAatgaccaacatccgcgtagc |
| RL-Cg.ddh:CO-fw | attcgtcatatgTACTAGAGAAAGAGGGGAAATAATCAatgaccaacatccgtgtagc |
| RL-Cg.ddh-rv | tagactgtcgacttagacgtcgcgtgcgatc |
| *Plasmid verification primers* | |
| ver-pET28-fw | ATGCGTCCGGCGTAGA |
| ver-pET28-rv | CTAGTTATTGCTCAGCGGT |
| ver-pMBI-fw | CGGGAAACGGTCATATAAG |
| ver-pMBI-rv | CTCTAGTAGAGAGCGTTCAC |

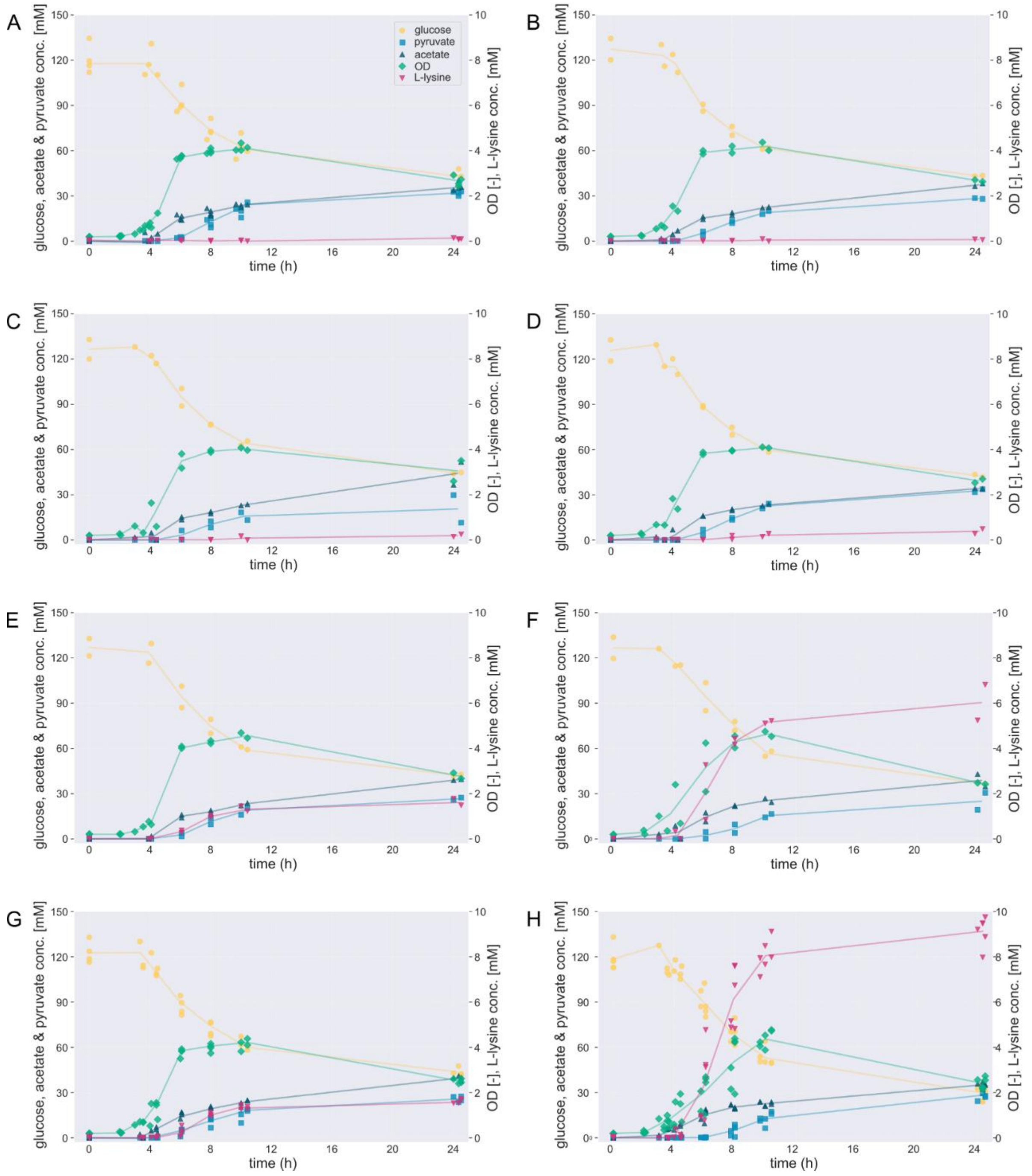


**Figure S1: Cultivation profiles of *V. natriegens Δdns* harboring (A) the pMBI empty plasmid, (B) the pMBI_Vn.LysC1, (C) the pMBI_Vn.LysC1:T353I, (D) the pMBI_Vn.LysC2, (E) the pMBI_Vn.dapA1, (F) the pMBI_Vn.dapA1:E84T, (G) the pMBI_Vn.LysC1:T353I_Vn.dapA1:E84T and (H) the pMBI_Vn.LysC2_Vn.dapA1:E84T vector during growth on glucose.** Time-course data showing biomass accumulation ($OD_{600\ nm}$), glucose consumption, and formation of pyruvate, acetate, and L-lysine. Scatter points show individual raw measurements. Lines represent tolerance-based mean values, where measurements within 0.5 h were grouped and averaged to visualize the central trend without altering the original data (25 mL S-limited VN medium with 120 mM glucose and 150 mM $NH_4Cl$, 1 mM IPTG, 37°C, 220 rpm).

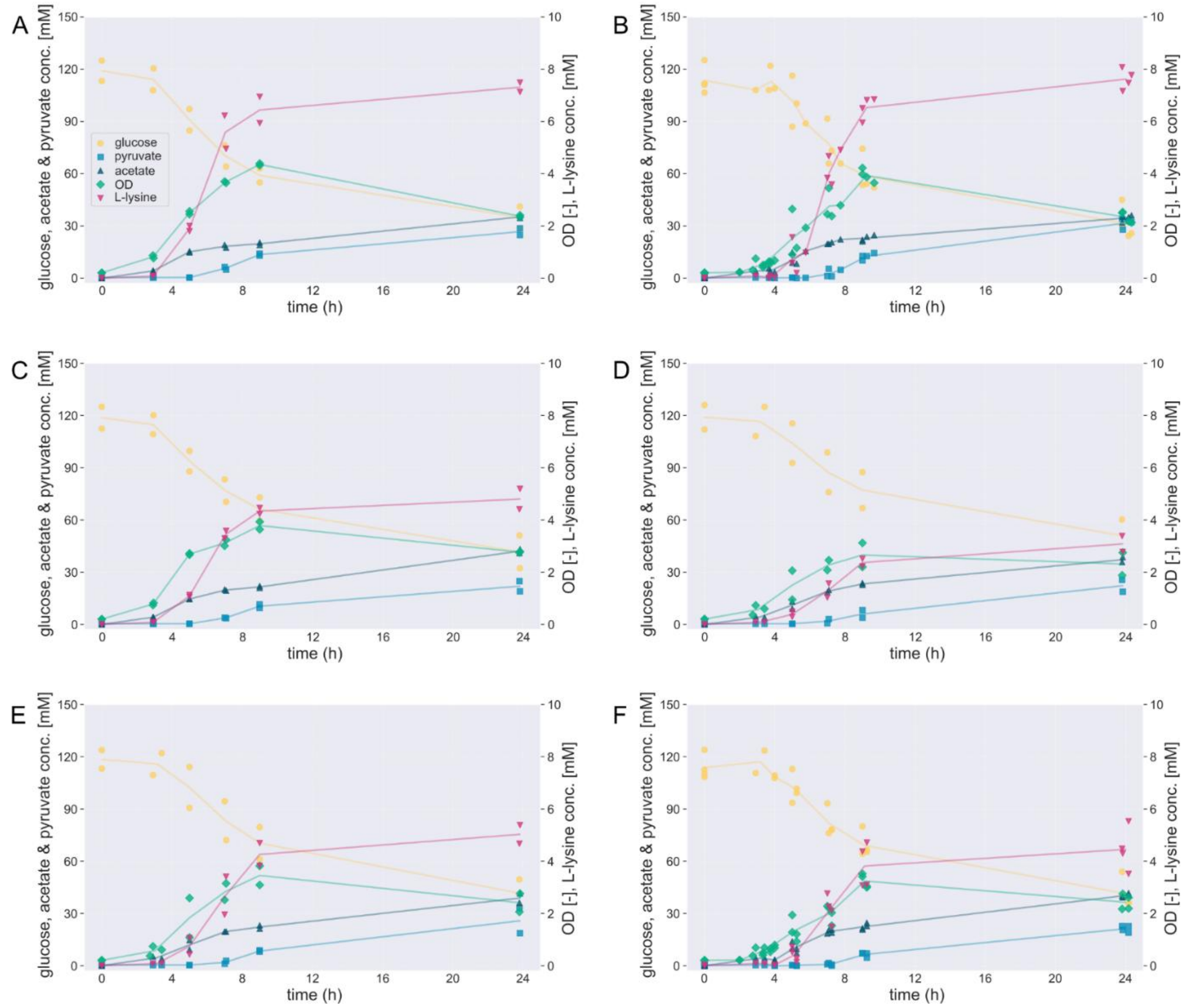


**Figure S2: Cultivation profiles of six *V. natriegens Δdns* strains during growth on glucose (A-F), each overexpressing additional L-lysine pathway enzymes from pMBI vectors. All plasmids share the same two upstream genes *(Vn.lysC2* and *Vn.dapA1:E84T)* , while the third gene and its RBS vary: (A) *Vn.dapD*-RBS32, (B) *Vn.dapD*-RBS64, (C) *Cg.ddh*-RBS32, (D) *Cg.ddh*-RBS64, (E) *Cg.ddh:CO*-RBS32, and (F) *Cg.ddh:CO*-RBS64.** Time-course data showing biomass accumulation ($OD_{600\ nm}$), glucose consumption, and formation of pyruvate, acetate, and L-lysine. Scatter points show individual raw measurements. Lines represent tolerance-based mean values, where measurements within 0.5 h were grouped and averaged to visualize the central trend without altering the original data (25 mL S-limited VN medium with 120 mM glucose and 150 mM $NH_4Cl$, 1 mM IPTG, 37°C, 220 rpm).

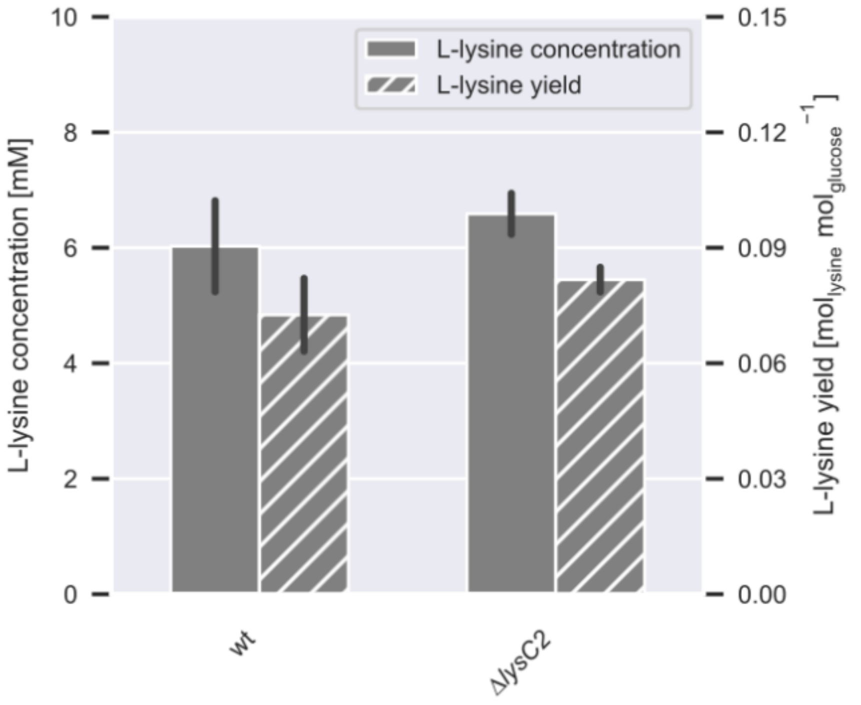


**Figure S3: Effect of *lysC2* deletion on L-lysine production in *V. natriegens* strains expressing the L-lysine insensitive Vn.dapA1:E84T variant.** L-lysine concentration after 24 h of cultivation and L-lysine yield of *V. natriegens* DSM759 *Δdns* (wt) and *V. natriegens* DSM759 *Δdns ΔlysC2* harboring the pMBI_Vn.dapA1:E84T vector during growth on glucose (25 mL S-limited VN medium with 20 g $L^{-1}$ glucose and 150 mM $NH_4Cl$, 1 mM IPTG, 37°C, 220 rpm).

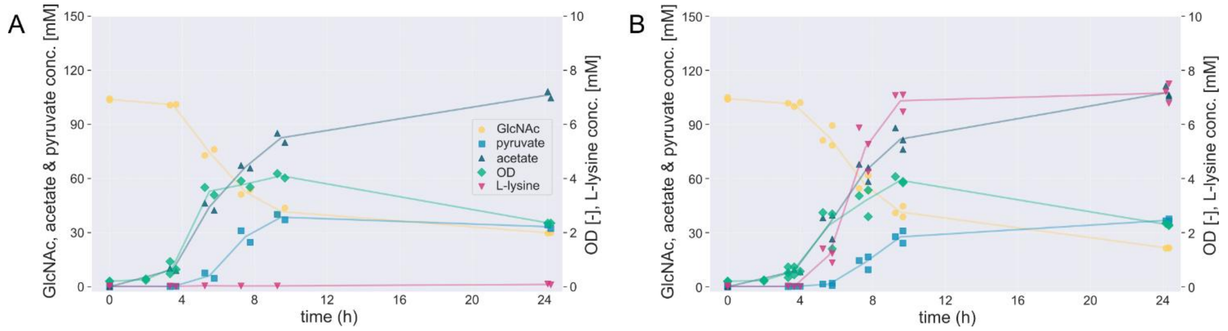


**Figure S4: Cultivation profiles of *V. natriegens Δdns* harboring (A) the pMBI empty plasmid or (B) the pMBI_Vn.LysC2_Vn.dapA1:E84T vector on GlcNAc as carbon source.** Time-course data showing biomass accumulation ($OD_{600\ nm}$), GlcNAc consumption, and formation of pyruvate, acetate, and L-lysine. Scatter points show individual raw measurements. Lines represent tolerance-based mean values, where measurements within 0.5 h were grouped and averaged to visualize the central trend without altering the original data (25 mL S-limited VN medium with 100 mM GlcNAc and 150 mM $NH_4Cl$, 1 mM IPTG, 37°C, 220 rpm, n≥2).
